# Microtubule depolymerization at kinetochores restricts anaphase spindle elongation

**DOI:** 10.1101/2024.08.30.610502

**Authors:** Geng-Yuan Chen, Changfeng Deng, David M. Chenoweth, Michael A. Lampson

## Abstract

Anaphase chromosome segregation depends on forces exerted by spindle microtubules. Current models propose two mechanisms of force generation: kinetochore microtubules (kMTs) depolymerize to pull chromosomes toward the spindle poles (anaphase A), while antiparallel microtubule sliding in the central spindle further separates the chromosomes by elongating the spindle (anaphase B). Experimental evidence in cells supports the sliding mechanism, but contributions of the depolymerization mechanism remain unclear. Here we show that kMT depolymerization limits spindle elongation rather than moving chromosomes apart. We developed a chemical optogenetic approach to recruit a microtubule depolymerase to kinetochores at anaphase onset, thereby increasing the rate of kMT depolymerization without perturbing earlier stages of mitosis. We find that increased depolymerization slows the velocity at which spindle poles move apart without changing kinetochore separation velocities. Our findings support a model in which kinetochores selectively couple to central spindle microtubules parallel to their kMTs, so that antiparallel sliding drives chromosome segregation while kMT depolymerization pulls poles inward.

## Introduction

During cell division, sister chromosomes segregate in anaphase to partition the two replicated genomes into daughter cells. Three mechanisms of force generation in anaphase have been described. First, kinetochore-microtubule (kMT) bundles depolymerize to pull chromosomes and spindle poles toward each other. Second, antiparallel microtubule sliding in the central spindle pushes the two half-spindles apart. Third, astral microtubules pull spindle poles apart. Connections between these different microtubule populations make it difficult to dissect the contributions of potential force-generating mechanisms to chromosome segregation.^1^

Current “chromosome-pulled” models propose that depolymerizing kMTs pull each chromosome toward its attached pole (anaphase A) (**Fig. 1a**).^2–4^ As chromosomes are pulled outward, antiparallel microtubule sliding pushes poles apart or pushes kMTs apart to elongate the spindle (anaphase B).^5,6^ Astral microtubules can also pull poles apart, but antiparallel sliding is thought to be the primary mechanism driving anaphase B in human cells.^6^ Molecular motors slide antiparallel microtubules apart *in vitro*, mimicking spindle elongation,^7–9^ and laser ablation of the central spindle between sister kinetochores in early anaphase slows kinetochore separation.^10,11^ Furthermore, manipulations that increase or decrease antiparallel sliding velocities affect both anaphase spindle elongation and chromosome segregation velocities.^12–18^ These findings demonstrate the importance of antiparallel sliding for chromosome segregation. In contrast, although depolymerizing microtubules can move tip-coupled objects (chromosomes or beads) in vitro,^3,4^ the contribution of kMT depolymerization to anaphase chromosome segregation remains unclear. Manipulations that slow kMT depolymerization in cells, by inhibiting microtubule depolymerase activity, have the expected effect of decreasing the velocity of kinetochores and poles moving toward each other (anaphase A). However, chromosome segregation velocities have not been reported in these conditions.^19–24^ In addition, depolymerases localize to microtubules beyond kMTs, so inhibition can affect multiple populations of spindle microtubules.^4^

**Figure 1.**
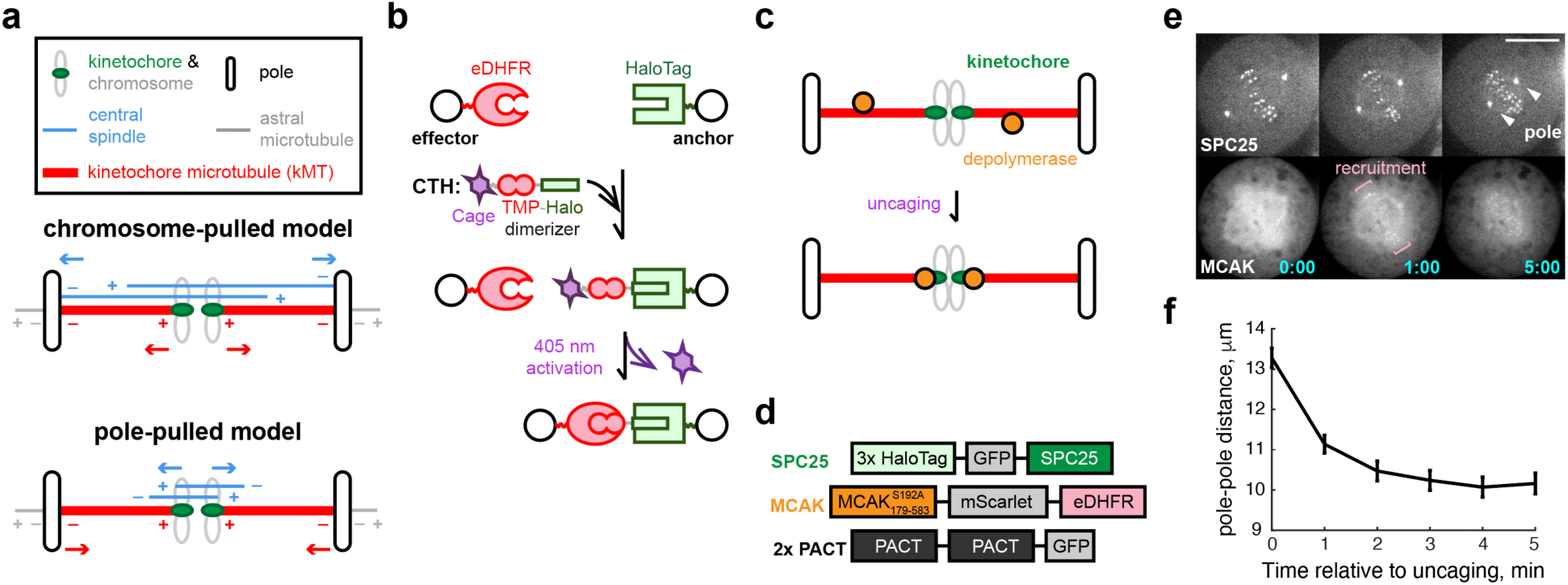
Induced kMT depolymerization shortens metaphase spindle. **(a)** Mechanical models for anaphase chromosome segregation. Polarity of antiparallel microtubules in the central spindle (blue), kMTs (red), and astral microtubules (gray) is indicated, with minus-ends (–) and plus-ends (+) facing toward and away from poles, respectively. In the chromosome-pulled model, antiparallel sliding in the central spindle pushes poles or kMTs apart (blue arrows), while kMT depolymerization pulls each chromosome outward toward its attached pole (red arrows). In the pole-pulled model, the central spindle couples to chromosomes, so antiparallel sliding directly moves sister chromosomes apart (blue arrows), and kMT depolymerization pulls poles inward toward the chromosomes (red arrows). **(b)** Schematic of light-inducible dimerization. The photocaged dimerizer (CTH) includes a Halo ligand (green) linked to the eDHFR ligand trimethoprim (TMP, red) and a photocage (purple). The dimerizer is activated by uncaging to make TMP accessible to eDHFR, recruiting the eDHFR-tagged effector to the Halo-tagged anchor. **(c)** Schematic of kMT shortening assay in metaphase. The microtubule depolymerase MCAK (orange) is recruited to kinetochores (green) after dimerizer uncaging at t=0 to enhance kMT depolymerization. **(d)** Constructs for kMT shortening assay. The minimal microtubule depolymerization domain (MCAK_179-583_) was used, with a mutation to prevent phosphorylation: S192A. SPC25 is the kinetochore anchor, and PACT labels spindle poles. **(e-f)** Metaphase kMT shortening assay for cells expressing Halo-GFP-SPC25, PACT-GFP, and MCAK-mScarlet-eDHFR. Representative images (e) show kinetochores, poles, and MCAK recruitment after dimerization at t=0. Pole-pole distances (f) were measured for each cell and averaged at every time point (n=29 cells pooled from four independent experiments). Time stamps, min:s. Scale bars, 10 µm. Error bars: mean ± SEM.

In an alternative model, the central spindle is coupled to chromosomes so that antiparallel sliding directly drives anaphase chromosome segregation.^25^ Neither kMT depolymerization (anaphase A) nor spindle elongation (anaphase B) is necessary for chromosome movements in this model. The spindle elongates because poles are attached to kMTs, so sliding moves poles outward, but chromosome movements are independent of spindle poles. We refer to this model as the “pole-pulled” model because kMT depolymerization at either kinetochores (plus-ends) or poles (minus-ends) pulls poles inward toward the chromosomes (**Fig. 1a**).^25^ Spindle elongation is thus a combination of outward forces on spindle poles from antiparallel sliding and astral microtubules and inward forces from kMT depolymerization and potentially central spindle depolymerization. For example, if kMT depolymerization is fast enough, chromosomes can segregate without spindle elongation as in *C. elegans* meiosis.^26^

To distinguish between the chromosome-pulled and pole-pulled models, we designed a chemical optogenetic assay to acutely increase microtubule depolymerization at kinetochores in anaphase in human cells. We show that increased kMT depolymerization restricts pole separation without affecting chromosome separation, consistent with the pole-pulled model. We propose a molecular model in which each kinetochore selectively binds to the sides of central spindle microtubules in parallel orientation to its kMTs. Antiparallel sliding moves central spindle microtubules of opposite orientations apart, along with their bound kinetochores. As kMTs depolymerize, poles are pulled inward toward kinetochores anchored to the central spindle. To test this model, we developed additional chemical optogenetic assays to manipulate spindle checkpoint signaling and antiparallel sliding in the central spindle. We inhibited the checkpoint to induce anaphase on monopolar spindles.^27^ Some sister kinetochore pairs localize near antiparallel microtubule bundles and move away from other in anaphase, consistent with antiparallel sliding driving their movements. We also recruited a microtubule-binding protein to the central spindle in anaphase to suppress antiparallel sliding. We find reduced kMT depolymerization, consistent with kinetochore linkage to the central spindle to constrain plus-end depolymerization. Overall, these results support our model in which kinetochores are mechanically coupled to central spindle microtubules oriented parallel to their kMTs, so that antiparallel sliding moves sister chromosomes apart while kMT depolymerization pulls poles inward.

## Results

### kMT depolymerization regulates spindle elongation

The chromosome-pulled model predicts that increasing kMT depolymerization in anaphase should increase kinetochore separation velocity, with no effect on pole separation velocity. Conversely, the pole-pulled model predicts decreased pole separation velocity as poles are pulled inward, with no effect on kinetochore separation velocity. To distinguish between these models, we designed an assay to increase the rate of microtubule depolymerization at kinetochores at anaphase onset. We recruited the motor domain of the microtubule depolymerase MCAK (Mitotic Centromere-Associated Kinesin; kinesin-13) to kinetochores using a photocaged small molecule that heterodimerizes HaloTag and *Escherichia coli* dihydrofolate reductase (eDHFR) fusion proteins.^28–30^ We fused HaloTag to the kinetochore protein SPC25 (the anchor) and fused eDHFR to the MCAK motor domain (the effector), so that uncaging leads to dimerization and MCAK recruitment to kinetochores (**Fig. 1b-d**). This chemical optogenetic approach provides temporal precision by light activation and molecular specificity by using genetically tagged constructs.^31^ By activating immediately after anaphase onset, our manipulations do not affect earlier stages of spindle formation and kinetochore attachment. As MCAK is active only at microtubule ends,^32^ we expect that recruitment of the motor domain to kinetochores increases depolymerization of kMTs but not other proximal microtubules in the central spindle.

We first tested the kMT depolymerization assay in metaphase spindles, in which cohesion prevents sister kinetochores from moving farther apart. We find that MCAK recruitment leads to rapid spindle shortening, as poles move inward toward the kinetochores within 2 min of uncaging (**Fig. 1e-f, Supplementary Video 1**). This result indicates that MCAK recruitment increases kMT depolymerization as expected, and that kinetochores remain coupled to their attached kMTs.

To determine the effects of increased kMT depolymerization in anaphase, we recruited MCAK to kinetochores within 1 min after sister kinetochores start to move apart, indicating anaphase onset (**Fig. 2a-b, Supplementary Video 2**). We measured three distances at each time point: between separating kinetochores, between poles, and between kinetochores and their attached pole (**Fig. 2c**). For each cell, we fit a slope to the earliest time window exhibiting a linear range for all distances to measure velocities of pole separation (V_PP_), kinetochore separation (V_KK_), and kMT depolymerization (V_PK_) (**Fig. 2d**). MCAK overexpression alone does not affect any of these velocities (**Supplementary Fig. 1**), and MCAK recruitment to kinetochores increases the rate of kMT depolymerization, as expected (V_PK_, from 0.52 ± 0.03 µm/min to 0.77 ± 0.04 µm/min, mean ± SEM, **Fig. 2e**). Pole separation velocity decreases (V_PP_, from 1.07 ± 0.08 µm/min to 0.40 ± 0.10 µm/min, mean ± SEM) with increased kMT depolymerization, but kinetochore separation velocity is not affected (**Fig. 2f-g**). The decrease in V_PP_ is double the increase in V_PK_, which represents only half of the spindle, indicating that poles are pulled inward toward the kinetochores by depolymerizing kMTs as in metaphase (**Fig. 1e-f**). We conclude that kMT depolymerization restricts spindle elongation rather than moving chromosomes, consistent with the pole-pulled model.

**Figure 2.**
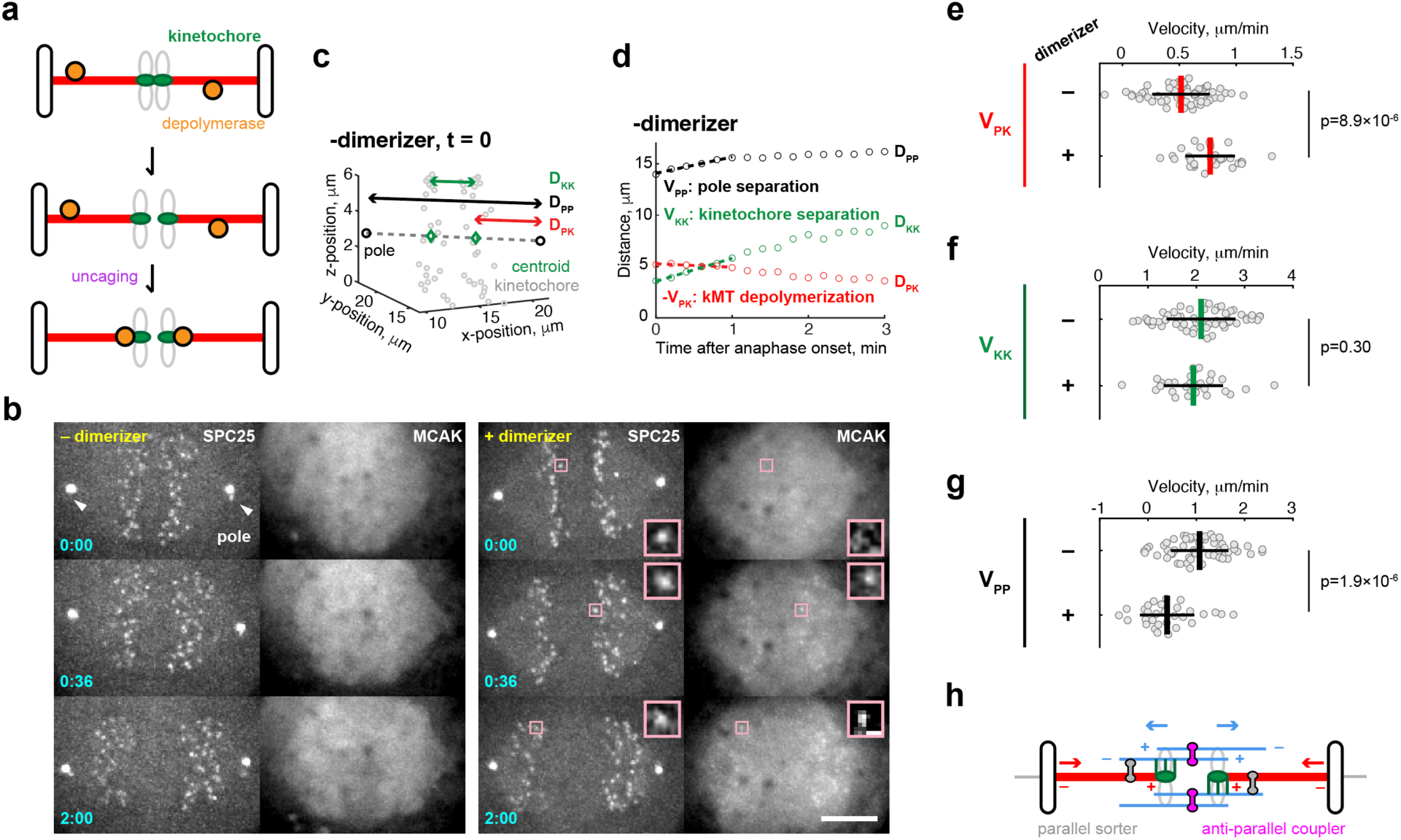
Enhanced kMT depolymerization slows pole separation in anaphase. **(a)** Schematic of kMT shortening assay in anaphase. The microtubule depolymerase MCAK (orange) is recruited to kinetochores (green) after dimerizer uncaging at t=0 at anaphase onset. **(b)** Representative images show cells expressing Halo-GFP-SPC25, MCAK-mScarlet-eDHFR, and PACT-GFP, with or without dimerization at t=0, immediately after sister kinetochores start to move apart. Contrast in the mScarlet channel was adjusted in each image to visualize MCAK recruitment. Insets for +dimerizer cells show kinetochores and MCAK recruitment at higher magnification with enhanced contrast. Time stamps, min:s. Scale bars, 5 µm or 500 nm in insets. **(c-d)** Analyses of kinetochore and pole motions. Example snapshot at t=0 (c) shows coordinates of poles (black) and kinetochores (gray circles). A pair of kinetochore centroids (green) represents the median positions of the two separating kinetochore masses projected to the mitotic axis (gray dashed lines, defined by the two poles). Three distances were calculated based on these coordinates: between kinetochore centroids (D_KK_, green), between poles (D_PP_, black), and between each kinetochore centroid and its adjacent pole (D_PK_, red). Example traces (d) show distances over time, with initial velocities (dashed lines) measured in the earliest time window exhibiting linear increases of all three distances. **(e-g)** Initial velocities were calculated for kMT depolymerization (e), kinetochore separation (f), and pole separation (g) (n=58 cells – dimerizer pooled from six independent experiments, 30 cells +dimerizer pooled from seven independent experiments). Each data point represents a single cell. Lines: mean ± SD. **(h)** The sort-and-grip model proposes that parallel sorters (gray dumbbells) position kMTs (red) adjacent to parallel microtubules in the central spindle (blue). Kinetochores (green) grip adjacent microtubules, which are in the correct orientation for antiparallel sliding (mediated by antiparallel couplers/motors, magenta dumbbells) to move sister kinetochores apart (blue arrows). Depolymerization at kMT plus-ends pulls poles inward toward kinetochores (red arrows) that are coupled to the central spindle.

To explain both outward forces on chromosomes and inward forces on poles, we propose a “sort and grip” model: each kinetochore sorts central spindle microtubules and selectively grips (binds tightly to the sides of) those in parallel orientation to the kMTs (**Fig. 2h**). As a result, antiparallel sliding moves kinetochores and their attached kMTs outward, while kMT depolymerization pulls poles inward relative to the central spindle where kinetochores are anchored. Forces exerted by kMT plus-end depolymerization rely on the well-established ability of kinetochores to maintain attachments to depolymerizing kMT plus-ends.^3^ Sorting can be mediated by molecular motors, crosslinkers, or branching microtubule arrays that sort microtubules for the same polarity,^33,34^ so that kinetochores selectively bind parallel central spindle microtubules based on proximity.

### Anti-poleward motions in monopolar anaphase

To test the sort-and-grip model, we designed two additional chemical optogenetic assays to manipulate spindle checkpoint signaling and antiparallel sliding in the central spindle. First, we initiated anaphase on monopolar spindles by inhibiting the spindle assembly checkpoint using conditional pharmacology. The sort-and-grip model predicts that kinetochore motions depend on the arrangement of kMTs and adjacent antiparallel bundles. For some sister kinetochore pairs, kMTs orient in opposite directions on a monopolar spindle (**Fig. 3a**).^35^ In this configuration, antiparallel sliding would move sister kinetochores away from each other rather than moving both sisters toward the pole. To induce anaphase, we used a photocaged inhibitor of the checkpoint kinase MPS1 (Monopolar Spindle 1) linked to the Halo ligand, which binds covalently to the Halo-tagged kinetochore anchor SPC25 (**Fig. 3b**).^36^ MPS1 is inhibited only after uncaging with a pulse of light. As the inhibitors are bound to SPC25 and do not diffuse between cells, local uncaging triggers anaphase in one cell at a time. This approach allows sequential live imaging of individual cells with high temporal resolution.

**Figure 3.**
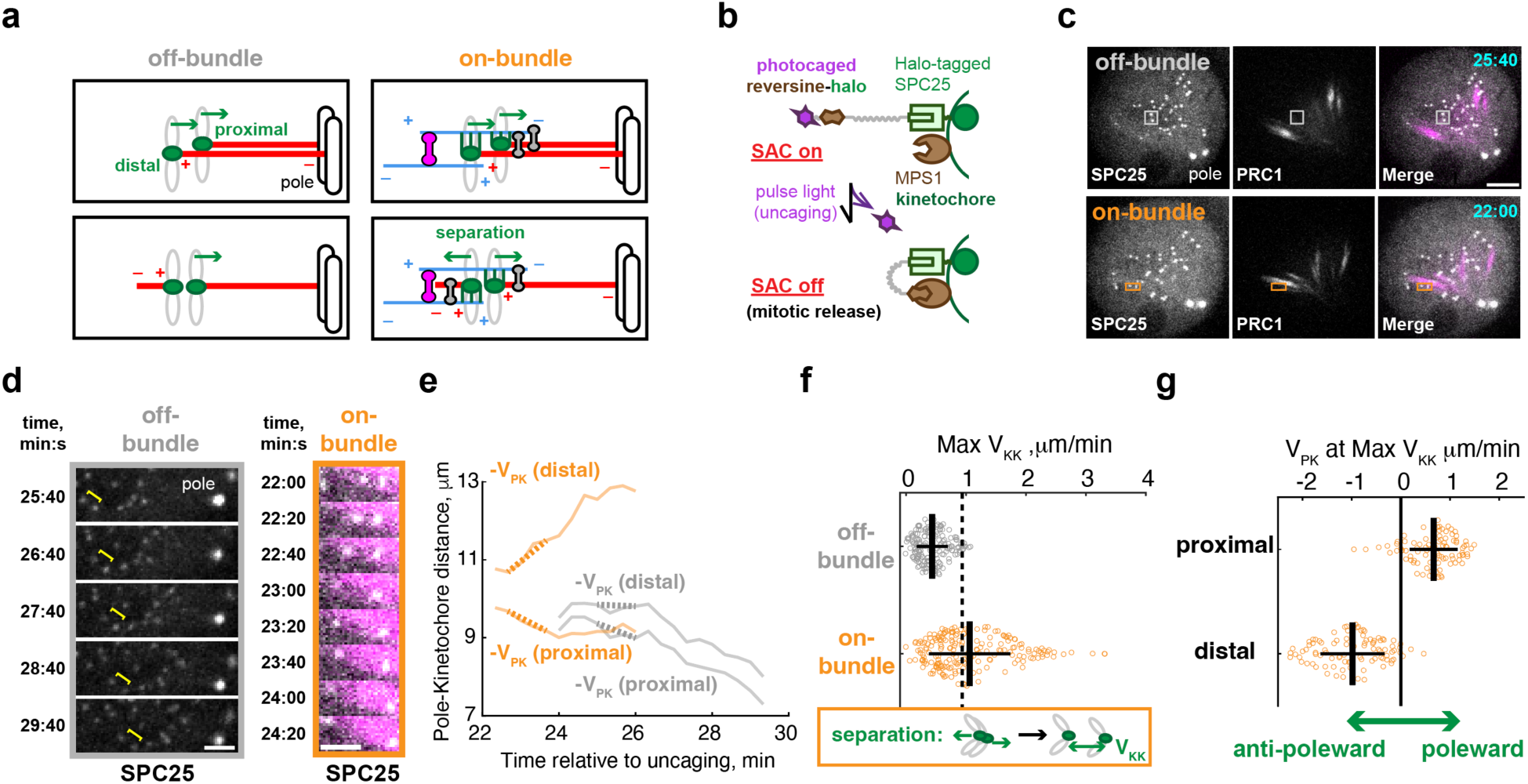
Anti-poleward kinetochore motions in monopolar anaphase. **(a)** Schematic of monopolar anaphase. Sister kinetochore pairs can localize away from or close to antiparallel bundles (off-bundle or on-bundle, respectively). Both kinetochores can attach to the pole (top panels), or the kMTs of the distal kinetochore can orient away from the pole (bottom panels). In the latter case, the sort-and-grip model predicts that antiparallel sliding can move the proximal and distal kinetochores toward and away from the pole, respectively, as kinetochores grip microtubules in parallel orientation to their kMTs (bottom right panel). Green arrows represent directions of kinetochore movement. **(b)** Schematic illustrates the conditional pharmacology approach for spindle checkpoint silencing in single cells. The MPS1 inhibitor (reversine, brown) is linked to a photocage (purple) and a Halo ligand (light green). The caged inhibitor localizes to kinetochores by binding to the Halo-tagged kinetochore protein SPC25 (green) and inhibits MPS1 (brown) only after uncaging. **(c-d)** Representative images (c) show HeLa cells expressing Halo-GFP-SPC25 (kinetochores), PACT-GFP (spindle poles), and 670nano3-PRC1 (antiparallel bundles), incubated with the kinesin-5 inhibitor STLC to create monopolar spindles. Time-lapse images (d) show representative off-bundle and on-bundle pairs, boxed in c. Yellow line: distance between the off-bundle sisters; time stamps (min:s): time after uncaging at t=0. **(e)** Individual kinetochore movements relative to the pole for the off-bundle (gray) and on-bundle (orange) sister pairs shown in d. Kinetochore velocity relative to the pole (V_PK_) was calculated as the displacement within a one-minute time window, with poleward motion defined as the positive direction. Dashed lines represent the maximal velocity of sister kinetochore separation (Max V_KK_). **(f)** Max V_KK_ plotted for off-bundle (n=168) and on-bundle (n=204) sister kinetochore pairs, from 23 cells pooled from seven independent experiments. Dashed line: upper bound for pairs that behave as off-bundle (95^th^ percentile of the Max V_KK_ distribution for off-bundle pairs). Some on-bundle pairs separate with high V_KK_ as shown in e (orange traces). **(g)** For each on-bundle pair with Max V_KK_ above the dashed line (n=93 pairs), indicating functional interactions with the bundle, V_PK_ is plotted for the proximal and distal sisters. Black lines: mean ± SD. Scale bars, 5 µm in c or 2 µm in d.

After MPS1 inhibition, we examined localization of the antiparallel microtubule crosslinker PRC1 (Protein Regulator of Cytokinesis 1), which marks antiparallel bundles.^14,37,38^ PRC1 signals gradually decrease in length and increase in intensity, indicating antiparallel sliding that decreases the overlap length in each bundle (**Supplementary Fig. 2**), as observed in the central spindle in normal bipolar anaphase^14,37,38^ and in reconstituted systems.^39–41^ We analyzed kinetochore motions separately for sister pairs either colocalized to (on-bundle) or away from (off-bundle) PRC1 signals (**Fig. 3c-f, Supplementary Videos 3-4**). Off-bundle pairs exhibit minimal movement of sisters away from each other, consistent with a lack of separation forces by antiparallel sliding. Conversely, some on-bundle sisters move apart, with the distal kinetochore of these pairs moving away from the pole (**Fig. 3g**). This result is consistent with the sort-and-grip model, in which antiparallel sliding drives chromosome movements depending on local spindle organization near kinetochores, independent of spindle poles. As monopolar spindles are generated by inhibiting kinesin-5 motors, antiparallel sliding is likely due to kinesin-4 (KIF4A) motors recruited to PRC1 bundles.^15^ Previous findings, using laser ablation to detach one kinetochore from its attached pole in a bipolar spindle, also show sister kinetochore separation consistent with the sort-and-grip model.^11^

### Suppressed central spindle sliding inhibits kMT depolymerization

In the sort-and-grip model, kMT plus ends are coupled to the central spindle via side-attached kinetochores. Without antiparallel sliding to move kinetochores outward, kMT depolymerization would require shortening the spindle by pulling poles inward, against outward forces exerted on poles by astral MTs. Thus, we predict that suppressed antiparallel sliding would reduce kMT depolymerization because both kMT plus and minus ends are constrained.

To suppress antiparallel sliding, we generated “brake” complexes in the central spindle. We fused HaloTag to full-length dimeric PRC1 (the anchor) and fused eDHFR to a monomeric PRC1 truncation mutant (PRC1M, the effector) that binds to microtubules without crosslinking activity on its own. Uncaging generates brake complexes by recruiting PRC1M to dimeric PRC1 bound to antiparallel bundles, which should strengthen PRC1 binding to microtubules and thus increase friction to suppress sliding within the bundle (**Fig. 4a-b**). Before uncaging, PRC1M binds to microtubules proportional to local microtubule density, with a subtle enrichment around the equator (**Fig. 4c, Supplementary Video 5**). This enrichment increases after activation, consistent with recruitment to PRC1-labeled antiparallel bundles in the central _spindle.14,37,38_

**Figure 4.**
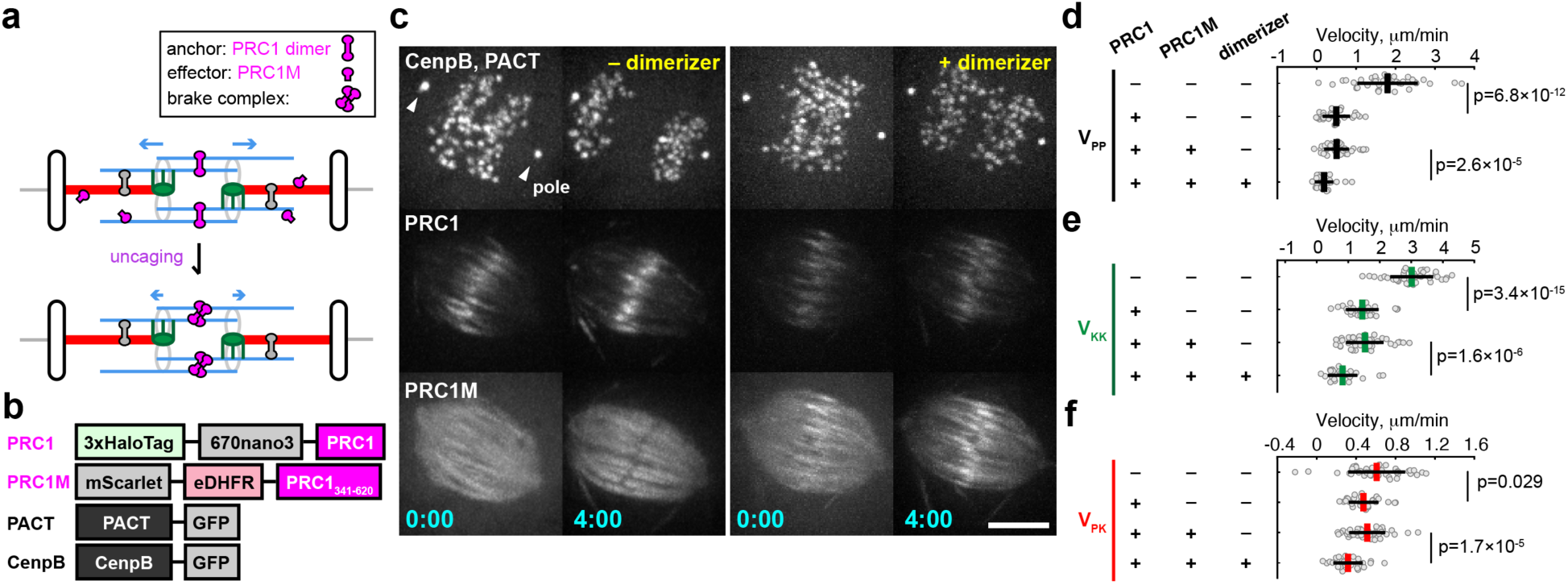
Suppressed bundle sliding inhibits kMT depolymerization. **(a)** Schematic of braking assay: monomeric PRC1 (PRC1M, magenta half-dumbbells) is recruited to dimeric PRC1 (magenta dumbbells) after uncaging at t=0, forming “brake” complexes to suppress antiparallel sliding (shown by reduced size of blue arrows). **(b)** Constructs for braking assay. PRC1M (PRC1_341-620_) is truncated to remove the N-terminus. PACT and CenpB label spindle poles and kinetochores, respectively. **(c)** Representative images of cells expressing CenpB-GFP, PACT-GFP, Halo-670nano3-PRC1, and mScarlet-eDHFR-PRC1M with or without dimerization at t=0, immediately after anaphase onset. Time stamps min:s; scale bars, 5 µm. **(d-f)** Initial velocity analyses for pole separation (d), kinetochore separation (e), and kMT depolymerization (f). From top to bottom in each panel: n=39 control cells (without Halo-670nano3-PRC1 or mScarlet-eDHFR-PRC1M), n=28 with Halo-670nano3-PRC1, n=42 with Halo-670nano3-PRC1 and mScarlet-eDHFR-PRC1M but no dimerization, n=27 with Halo-670nano3-PRC1 and mScarlet-eDHFR-PRC1M and dimerization. All cells express CenpB-GFP and PACT-GFP. Cells were pooled from three independent experiments; each data point represents a single cell. Lines: mean ± SD.

PRC1 itself acts as a molecular brake, as it restricts microtubule sliding *in vitro,*^39–41^ and its depletion in cells increases the separation velocity of the two half-spindles.^14,42^ Thus, we predict increased friction and suppressed antiparallel sliding due to PRC1 overexpression alone, followed by a further increase in friction after uncaging and PRC1M recruitment at anaphase onset. We measured kinetochore and pole velocities and kMT depolymerization as in Fig. 2c-d. We find that PRC1 overexpression decreases pole and kinetochore separation velocities, proportional to PRC1 levels on spindles, and PRC1M recruitment further decreases these velocities (**Fig. 4d-e, Supplementary Fig. 3a-c, Supplementary Video 6**). These results indicate progressively greater suppression of antiparallel sliding as expected. The rate of kMT depolymerization decreases as antiparallel sliding is suppressed, with the slowest rate observed in the presence of both PRC1 overexpression and PRC1M recruitment (**Fig. 4f, Supplementary Fig. 3d**). This result is consistent with the sort-and-grip model. The plus-ends of kMTs are coupled to the central spindle, while the minus-ends are anchored at the poles and constrained by astral MTs, so kMTs are not free to depolymerize.

## Discussion

Using chemical tools to manipulate spindle organization and specific microtubule populations within the spindle, our experiments support a model in which antiparallel sliding directly drives anaphase chromosome segregation while kMT depolymerization limits spindle elongation. We first show that enhanced kMT depolymerization slows pole separation without increasing kinetochore separation (**Fig. 2**). Based on these findings, we propose a sort-and-grip model (**Fig. 5**): parallel microtubule sorters position kinetochores close to central spindle microtubules oriented parallel to the kMTs, so that kinetochores selectively grip these central spindle microtubules upon anaphase onset. If sorting is established before anaphase onset, then sorters can be absent in anaphase or allow sliding of kMTs relative to the central spindle. Kinetochores grip tightly in anaphase, however, without sliding on the central spindle, which has two key outcomes. First, antiparallel sliding in the central spindle moves sister kinetochores apart together with their attached kMTs. Second, kMT plus-end depolymerization pulls kMTs and poles inward toward kinetochores that are tightly coupled to the central spindle, against outward movement of kMTs driven by anti-parallel sliding. Depolymerization of kMT minus-ends also pulls poles inward in this model. Our results from two additional assays support the sort-and-grip model. First, sister kinetochores can move apart on monopolar spindles, consistent with antiparallel sliding moving sorted microtubules together with their gripped kinetochores (**Fig. 3**). Second, suppressed antiparallel sliding slows kMT depolymerization, indicating that kinetochores couple to the central spindle (**Fig. 4**).

**Figure 5.**
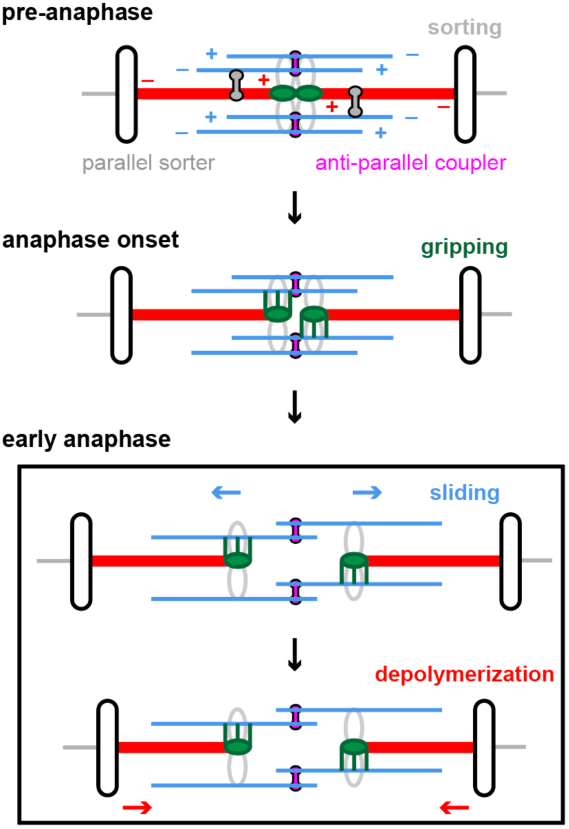
Proposed sort-and-grip processes. Before anaphase onset, microtubule sorting mechanisms position kMTs (red) close to parallel microtubules in the central spindle (blue) by parallel sorters (gray dumbbells). Upon anaphase onset, kinetochores (green) grip adjacent microtubules, which are correctly oriented for antiparallel sliding to move sister kinetochores apart. Parallel sorters are not needed after anaphase onset and are not shown for simplicity. Two concurrent processes drive pole and kinetochore movements in anaphase: antiparallel sliding directly separates gripped kinetochores, while microtubule depolymerization at kinetochores pulls poles inward toward kinetochores. The processes are shown sequentially for clarity but can occur simultaneously.

Velocity measurements in multiple model systems are consistent with the sort-and-grip model. In human cells, *C. elegans* oocytes and embryos, and *Xenopus* egg extracts, antiparallel sliding velocity within the central spindle matches kinetochore velocity in anaphase (**Supplementary Fig. 4a**)^10,43^, suggesting that antiparallel sliding moves chromosomes without requiring a contribution from kMT depolymerization. Sliding velocity also correlates with poleward movements of kMTs.^11,15^ In addition, slowing kMT depolymerization, by depleting microtubule depolymerases or severing enzymes in human cells or adding non-degradable cyclin B to *Xenopus* egg extracts, increases spindle elongation velocity.^20,24,44^ These results are consistent with kMT depolymerization pulling poles inward and with our findings that increasing kMT plus-end depolymerization slows spindle elongation (**Fig. 2**).

Microtubule severing experiments in multiple systems are also consistent with the sort-and-grip model. Cutting the central spindle early in anaphase transiently halts kinetochore separation in human cells and *C. elegans* oocytes and embyros,^10^ whereas kinetochores continue to separate after ablating centrosomes or microtubules midway between kinetochores and their attached poles in human, *C. elegans*, rat-kangaroo, newt, grasshopper, and crane-fly cells.^10,11,45–50^ These findings indicate that antiparallel sliding directly moves chromosomes. Structurally, central spindle microtubules are close to kinetochores and their attached kMTs in high-resolution optical and EM images,^10,51,52^ and kMT perturbations by microneedle manipulation or microtubule severing suggest crosslinking to the central spindle near chromosomes.^11,53–55^ Kinetochores can thus directly bind to the central spindle or interact with microtubule-associated proteins on the central spindle to make the coupling robust.

To move chromosomes toward their attached poles correctly, the sort-and-grip model relies on kinetochores attaching selectively to the sides of central spindle microtubules oriented parallel to their kMTs (**Fig. 5**). This selectivity can be mediated by microtubule sorting mechanisms acting earlier in mitosis that position kMTs adjacent to parallel microtubules in the central spindle. In prometaphase, short microtubules bind kinetochores at their plus-ends, while the minus-ends interact with antiparallel bundles in the central spindle.^56^ Minus-end directed motors sort these nascent kMTs and establish parallel coupling to central spindle microtubules to biorient sister kinetochores (**Supplementary Fig. 4b**).^57^ Alternatively, minus-end directed motors also sort newly-grown microtubules in the central spindle poleward through kMTs (**Supplementary Fig. 4c**).^58^ Furthermore, parallel microtubule arrays can assemble by growing microtubules from existing microtubule templates.^33,59–61^ Microtubules nucleated from sister kMTs can thus grow toward the equator to generate a “bridging fiber” by antiparallel crosslinking (**Supplementary Fig. 4d**).^11^ We propose that parallel microtubules sorted before anaphase persist so that kinetochores couple to these microtubules and move by antiparallel sliding in anaphase. Inhibition or depletion of potential candidates for these sorting mechanisms (e.g. augmin,^62^ NuMA,^63^ ZW10,^64^ and kinesin-14^65^) commonly causes anaphase chromosome segregation errors, which can arise from incorrect kinetochore coupling to microtubules antiparallel to their kMTs.

Restricting spindle elongation may be important for later stages of division, when excessive spindle elongation is associated with midzone buckling and defocused cytokinetic furrows.^14,37,38,66–70^ In addition, spindle poles act as hubs for numerous biochemical and mechanical cues,^71–73^ so restricted spindle elongation may regulate pole communication with spindle midbodies or the cytokinetic furrow. Hence, telophase biology likely requires coordination between pole separation and chromosome separation velocities via depolymerizing kMTs.

## Methods

### Synthesis, characterization, and storage of chemical optogenetic probes

The photoactivatable dimerizer (CTH)^29,30^ and photoactivatable localized MPS1 inhibitor (CRH)^36^ were prepared as previously described. Both compounds were dissolved in DMSO at 20 mM and stored as 1 μL aliquots in amber plastic microcentrifuge tubes at −80 °C.

### Small molecule inhibitors

All stocks were dissolved in DMSO and stored at −20 °C. Stock concentrations include 20 mM S-Trityl-L-Cysteine (STLC, Eg5 inhibitor; Sigma Aldrich) and 10 mM Verapamil (Tocris). Fresh aliquots were used for all experiments.

### Plasmids

The pEM705 backbone with a CAG promoter was used to clone all plasmids for anchors and effectors.^74^ The kinetochore anchor SPC25, fused to three tandem copies of HaloTag and an eGFP on its N-terminus (3×Halo-GFP-SPC25), was used as previously described.^29,30^ To manipulate central spindle mechanics, PRC1 was used as an anchor, fused to 3× HaloTag with the far-red fluorescent reporter, miRFP670nano3 (hereafter denoted 670nano3), on its N-terminus (3×Halo-670nano3-PRC1).

All effectors were designed based on the construct with the red fluorescent protein mScarlet fused to the N terminus of *Escherichia coli* dihydrofolate reductase DHFR (mScarlet-eDHFR). For targeted microtubule depolymerization, the phospho-resistant motor domain of human MCAK (MCAK_179-583;S192A_) was fused to the N terminus of mScarlet-eDHFR (MCAK-mScarlet-eDHFR).^30,75^ To suppress microtubule sliding, a monomeric PRC1 mutant (PRC1M, PRC1_341-620_)^76^ was fused to the C terminus of mScarlet-eDHFR to make mScarlet-eDHFR-PRC1M.

To label spindle poles, a single copy or two tandem copies of the PACT (Pericentrin and AKAP Centrosome Targeting)^77^ domain was fused to eGFP (PACT-GFP or 2×PACT-GFP) in the pcDNA3.1 backbone containing cytomegalovirus and T7 promoters. To label the central spindle, full-length PRC1 was fused to two tandem copies of 670nano3 (2×670nano3-PRC1), using pEM705 as the backbone.

### Cell cultures and transfection

HeLa RMCE (recombination-mediated cassette exchange) acceptor cells were passaged at 37°C in the growth medium containing DMEM (Gibco) supplemented with 10% Fetal Bovine Serum (Clontech) and 1% penicillin/streptomycin (Life Technologies), within 5% CO_2_ atmosphere and humidity. For assays, cells were cultured on coverslips (Bellco) coated by poly-D-lysine (Sigma-Aldrich) for at least 16 h, and then transfected with the desired constructs using Lipofectamine 3000 (Invitrogen) in Opti-MEM (Gibco) for at least 24 h.

Manipulations of mechanics and biochemistry at kinetochores were conducted with cells stably expressing 3×Halo-GFP-SPC25, as previously described.^29,30^ For the monopolar anaphase assay, cells were transfected with 0.5-1 µg 2×670nano3-PRC1 and 250 ng PACT-GFP to mark the central spindle and spindle poles, respectively. For kMT depolymerization, cells were transfected with 1-1.5 µg MCAK-mScarlet-eDHFR and 250 ng 2×PACT-GFP.

Manipulations of central spindle mechanics were conducted with cells transiently transfected with 0.6 µg Scarlet-eDHFR-PRC1M and 0.6 µg 3×Halo-670nano3-PRC1, together with 100 ng CenpB-GFP and 250 ng PACT-GFP for labeling centromeres and poles, respectively.

### Photoactivation and dimerization

For recruitment experiments, cells were placed in a magnetic chamber (Chamlide CM-S22-1; LCI) and incubated with growth medium and CTH for 1 h. Unbound CTH was washed out by incubating for 20 min in imaging medium containing L15 without phenol red (Gibco) (imaging medium) plus 10% FBS and 1% penicillin/streptomycin. The medium was then replaced with fresh imaging medium, and the chamber was moved to a stage in a 37°C environmental chamber (PeCon). For dimerization experiments, cells were imaged live to determine the time of anaphase onset. CTH was activated within 1 min after sister kinetochores started to move apart, using a 405-nm laser (CrystaLaser LC) controlled by iLas2 software (Roper Scientific) installed on MetaMorph (Molecular Devices), as previously described.^30^ Activation was performed over the entire cell at 25% laser power with 5 repetitions. To induce anaphase in monopolar cells, cells were incubated with 10 µM Verapamil (an efflux pump inhibitor) and 20 µM STLC in growth medium for 2 h, then 20 µM CRH was added for 1 h. Unbound CRH was washed out by incubating in imaging medium with STLC for 30 min. CRH was uncaged by a 5-second light pulse using a mercury arc lamp filtered at 385/11 nm, and imaging started 15-20 min later. Procedures with light-sensitive molecules were performed under red light in the room.

### Image acquisition

Live cells at 37 °C were imaged with a confocal microscope (DM4000; Leica), equipped with a 100× 1.4 NA oil immersion objective (Leica), an XY Piezo-Z stage (Applied Scientific Instrumentation), a CSU10 spinning disk (Yokogawa), an electron multiplier charge-coupled device camera (ImageEM; Hamamatsu Photonics), and a 4-line laser module (405 nm, 488 nm, 561 nm, and 639 nm; Vortran Stradus VersaLase 4), as previously described.^30^ Images were acquired with a 10-60s time interval, with 0.5-2 µm spacing for z-stacks covering 6-15 µm total. For dimerization experiments, cells were activated after image acquisition at t=0. For monopolar anaphase assay, images were acquired 15 minutes after uncaging.

### Data analyses and statistical tests

For monopolar anaphase cells, both kinetochores and poles were tracked by the Fiji::MTrackJ plug-in. In each frame, the distances of two sisters relative to the pole were calculated (D_PK_). The distance difference between the two sisters (ΔD_PK_) represents the inter-kinetochore distance along the axis between the pole and kinetochores (D_KK_). For each trajectory, the maximum kinetochore separation velocity (max V_KK_) represents the maximal displacement of inter-kinetochore distance (D_KK_) across all possible one-minute time windows. A cutoff was defined as the 95% percentile of max V_KK_ of off-bundle sister kinetochore pairs.

For bipolar anaphase cells, the coordinates of kinetochores and spindle poles were measured by the Fiji::TrackMate and Manual Tracking plug-ins, respectively. Kinetochore coordinates were projected to the mitotic axis defined by poles, and then grouped based on their closest pole. In each frame, the median position of each group represents the centroid of projected kinetochores. Initial velocities were defined by the time window exhibiting linear displacement between the two poles (V_PP_), between the two kinetochore centroids (V_KK_), and between a kinetochore centroid and its corresponding pole (-V_PK_). Cells were included in the analysis only if they exhibit constant kinetochore separation velocity at early time points, representing complete loss of cohesion and separation of sister chromosomes. To measure PRC1 intensity on a bipolar spindle, image stacks were mean-projected along z. Spindle-bound PRC1 signal was defined by the intensity within the minimal circle encompassing both spindle poles, with cytosolic intensity subtracted as background, summed for each cell. For each set of experiments, PRC1 intensities across different perturbations were normalized relative to the mean intensity of cells expressing 3×Halo-670nano3-PRC1.

Data from measurements were analyzed by MATLAB. Two-tailed Student’s t-test was used to compare measurements between groups.

## Acknowledgments

We thank the Philly ChromoClub and the Penn Center for Genome Integrity for discussions. This work was supported by the NIH (GM122475 and P01-CA265794) and the Basser Center for BRCA.

## Author contributions

GYC designed and conducted cell biology experiments and wrote the manuscript. CD and DMC synthesized and characterized chemical optogenetic probes. MAL designed experiments and edited the manuscript.

## Materials & Correspondence

Cell lines, chemicals, and protocols are available upon request. The corresponding authors adhere to the NIH Grants Policy and Sharing of Unique Research Resources. Correspondence and requests for materials should be addressed to MAL.

## Supplementary Information

is available for this paper.

## Data availability

The datasets generated and analyzed during the current study are available from the corresponding author on reasonable request without restriction.

---

**Supplementary Figure 1.**
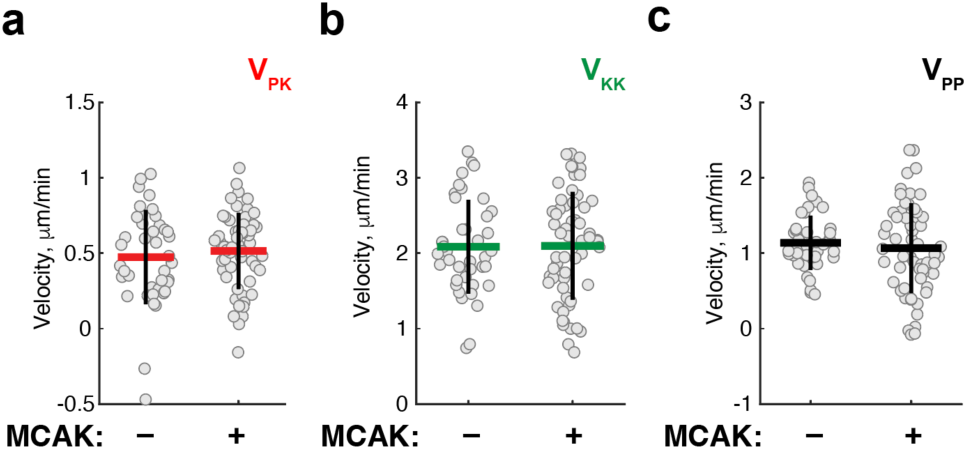
MCAK overexpression does not affect motions of chromosomes and poles. Velocities of kMT depolymerization (a), kinetochore separation (b), and pole separation (c) were measured for cells expressing mScarlet-eDHFR (–MCAK, n=40 cells), or MCAK-mScarlet-eDHFR (+MCAK, n=58 cells) as in Fig. 2c-d. For +MCAK cells, data were replotted from –dimerizer cells in Fig. 2e-g. Each data point represents a single cell. Lines: mean ± SD.

**Supplementary Figure 2.**
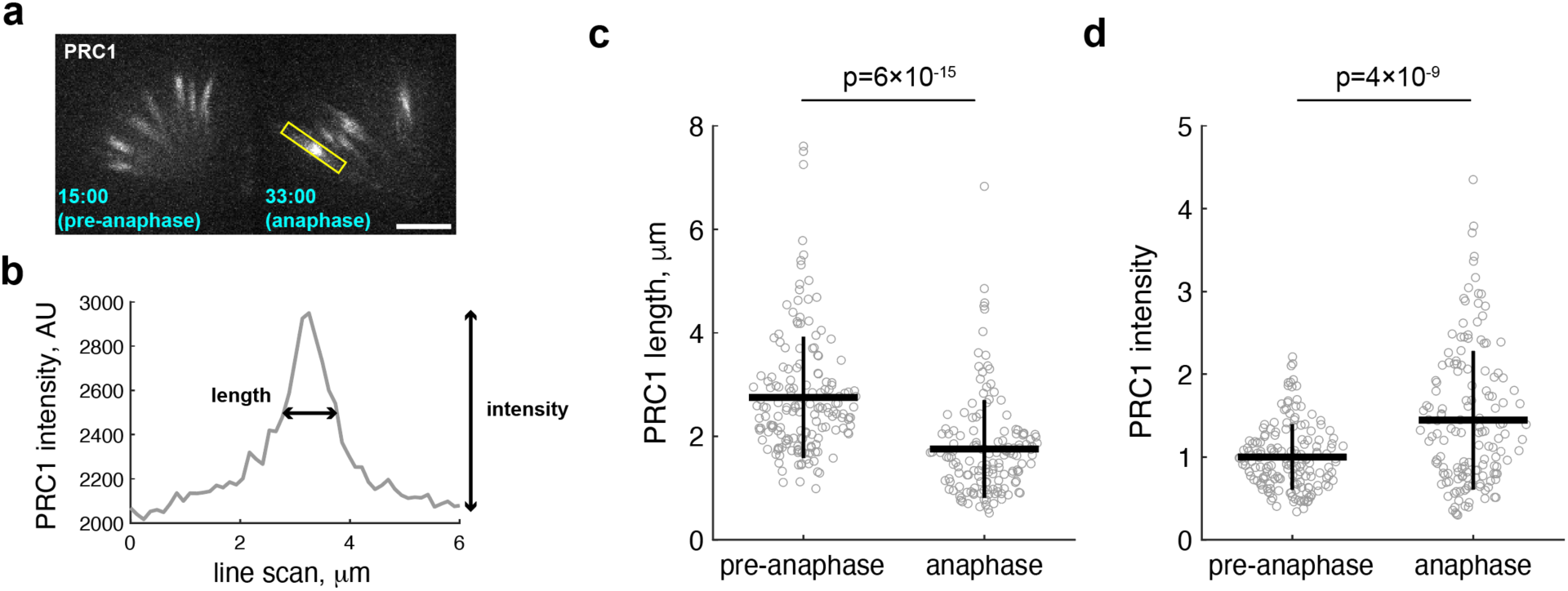
PRC1 morphology in monopolar spindles. **(a)** Representative images of PRC1 bundles before (left) or after (right) anaphase onset. Time stamps indicate the time after uncaging. Scale bars, 5 µm. **(b)** An example of PRC1 intensity analyzed by line scan (box in a). Bundle length and intensity are defined by the full width at half-maximum and the peak height, respectively. **(c-d)** Analyses for bundle length (c) and intensity (d). Each dot represents a PRC1 bundle (n = 162 for pre-anaphase bundles and 148 for anaphase bundles from 23 cells). Lines: mean ± SD.

**Supplementary Figure 3.**
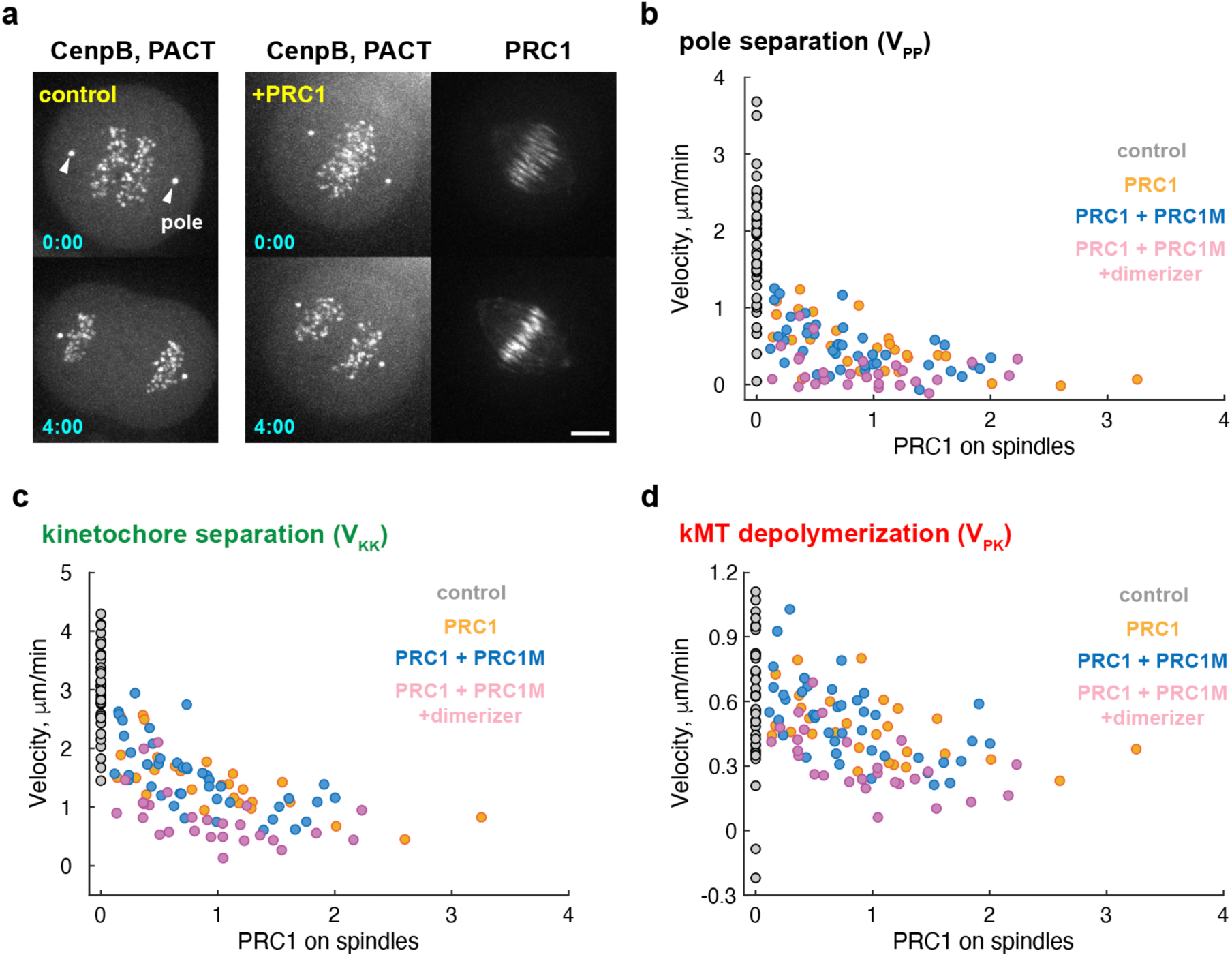
PRC1 overexpression and formation of brake complexes inhibit kMT depolymerization. **(a)** Representative images of cells expressing CenpB- GFP and PACT-GFP, with or without expression of Halo-670nano3-PRC1. Time stamps (min:s) indicate time after first observing sister kinetochore separation at t=0. Scale bars, 5 µm. **(b-d)** Dependence of pole separation (b), kinetochore separation (c), and kMT depolymerization (d) velocities on spindle-bound Halo-670nano3-PRC1. Colored data points show control cells without Halo-670nano3-PRC1 or mScarlet-eDHFR- PRC1M (gray), cells expressing Halo-670nano3-PRC1 (orange), cells expressing Halo- 670nano3-PRC1 and mScarlet-eDHFR-PRC1M without dimerization (blue), and cells expressing Halo-670nano3-PRC1 and mScarlet-eDHFR-PRC1M with dimerization (pink). All cells express CenpB-GFP and PACT-GFP. Each data point represents a single cell.

**Supplementary Figure 4.**
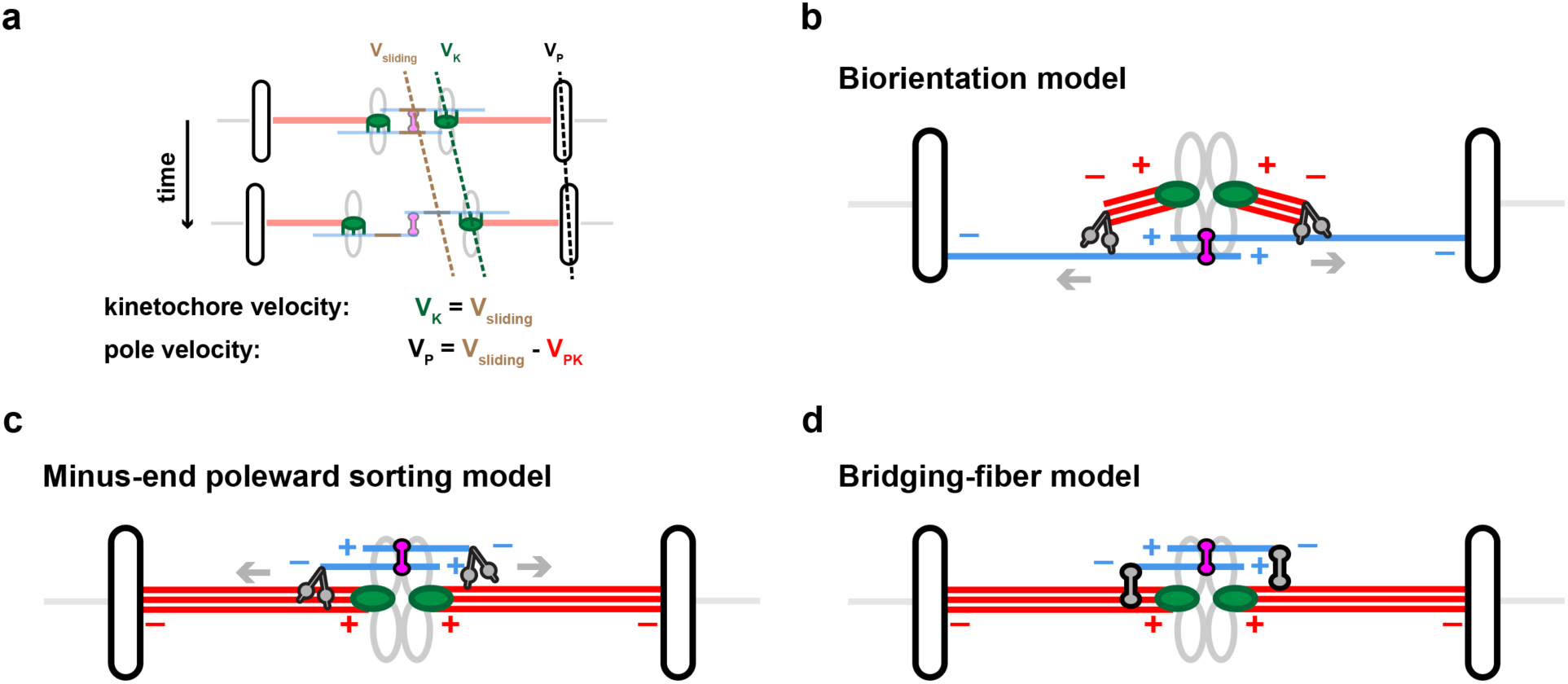
Velocity relationships predicted by the sort-and-grip model and proposed mechanisms of parallel sorting. (**a**) Schematic shows kinetochore velocity (V_K_), pole velocity (V_P_), and antiparallel sliding in the central spindle (V_sliding_). Kinetochore and pole velocities are half of the velocities of kinetochore separation (V_KK_) and pole separation (V_PP_), respectively, as measured in Figs. 2 and 4. Kinetochores, poles, and fiducial marks that label antiparallel sliding are shown in green, black, and brown, respectively. In the sort-and-grip model, kinetochore velocity matches sliding velocity, and pole velocity is sliding velocity (moving poles outward) subtracting kMT depolymerization from either end (V_PK_, moving poles inward). Manipulations of kMT depolymerization affect pole velocity but not kinetochore velocity in this model. (**b**) Kinetochore-nucleated kMT fragments (red) have their plus-ends at kinetochores. Minus-end directed motors (gray) at the minus-ends of these short kMTs preferentially guide kMT growth toward the minus ends of parallel microtubules in the central spindle (blue) to biorient sister kinetochores. (**c**) Minus-end directed motors (gray) guide the minus ends of newly-grown microtubules in the central spindle toward spindle poles through kMTs. (**d**) A fraction of the central spindle is grown by microtubules branching from kMTs (gray). These branching microtubules form antiparallel bundles (bridging fibers) between the two sister kinetochores, bound by antiparallel couplers.

**Supplementary Video 1.** Max-intensity projection over 1 µm for the cell shown in Fig. 1e. Time stamps (min:s) indicate time after uncaging at t=0. White arrows: spindle pole; scale bar: 10 µm.

**Supplementary Video 2.** Max-intensity projection over 4 µm for the cells shown in Fig. 2b, with (right) or without (left) dimerization. Time stamps (min:s) indicate time after uncaging at t=0. White arrows: spindle pole; scale bar: 5 µm.

**Supplementary Video 3.** Max-intensity projection over 2 µm showing the off-bundle sister kinetochore pair (yellow arrow) indicated by the gray box in Fig. 3c. Time stamps (min:s) indicate time after uncaging at t=0. Scale bar: 5 µm.

**Supplementary Video 4.** Max-intensity projection over 2 µm showing the on-bundle sister kinetochore pair (yellow arrow) indicated by the orange box in Fig. 3c. Time stamps (min:s) indicate time after uncaging at t=0. Scale bar: 5 µm.

**Supplementary Video 5.** Max-intensity projection over the whole cell for the cells shown in Fig. 4c. Left: –dimerizer; Right: +dimerizer. Time stamps (min:s) indicate time after uncaging at t=0. White arrows: spindle pole; scale bar: 5 µm.

**Supplementary Video 6.** Max-intensity projection over the whole cell for the cells shown in Supplementary Fig. 3a, with (right) or without (left) expression of Halo-670nano3-PRC1. Time stamps (min:s) indicate time after first observing sister kinetochore separation at t=0. White arrows: spindle pole; scale bar: 5 µm.

